# Bilateral asymmetry favours the left hemimandibular side in companion rabbits

**DOI:** 10.1101/351320

**Authors:** Pere M. Parés-Casanova, Maria Cabello, Baba Gana Gambo, Michael Oluwale Samuel, James Olukayode Olopade

## Abstract

Directional asymmetry (DA) appears when the left and right body sides differ consistently from each other. This asymmetry is characterized as a type of natural asymmetry typical of the population’s biology, which can be derived both from genetic inheritance, as of the functional importance acquired by certain features with respect to the environment in which they develop. We report here preliminary findings that for the first time to quantify size asymmetry mandibles of companion rabbits. A total sample of 64 companion rabbits from the same farm were studied by means of geometric morphometric techniques. The mandible morphology was described by a set of 18 landmarks and semilandmarks on the lateral aspect. Tests showed paired differences with variations in distributions, demonstrating a DA in favour of left mandibular side. The detected unilaterality could be interpreted as a manifestation of lateralized masticatory activity. This is a first time report to quantify size asymmetry in this species.

## Introduction

The left-right symmetry of hemimandibles corresponds to matched symmetry, where two separate objects exist as mirror images of each other. The object symmetry, in contrast, is referred to in a single structure is identical according to a given or selected plane, such as mid-sagittal plane. Like many skeletal structures, the mandible is generally assumed to be bilaterally symmetrical. The disturbances in symmetry and an occurrence of asymmetry within data might be an indicator of individual or population-related developmental stress, shed light on pathological conditions or indicate a relation between structurally or functionally interacting elements (1).

Developmental instability (DI) arises from genetic or environmental stressors that disturb the normal developmental pathways of different continuous characteristics, producing developmental instability (1). This is commonly measured as fluctuating asymmetry (FA) (2). FA is the variance in subtle differences between the left and right sides in bilaterally symmetrical organisms or parts of organisms, and is considered a measure of how well an individual can buffer its development against stressing factors and the resulting perturbations during development (3). Conversely, directional asymmetry (DA) appears when the left and right body sides differ consistently from each other(3) (2). Its expression is mediated by a left-right axis conveying distinct positional identities for developing structures on either body side (3).This asymmetry is characterized as a type of natural asymmetry typical of the population’s biology, which can be derived both from genetic inheritance, as of the functional importance acquired by certain features with respect to the environment in which they develop. Finally, the antisymmetry (AS) corresponds to a systematic deviation from symmetry, but in this case the side that is larger varies at random among individuals (2).

FA can be separated from two other forms of bilateral asymmetry based on the distribution of signed asymmetry values in the population. For a trait showing ‘ideal’ FA, the right-left differences are normally distributed around a mean of zero. DA is characterized by a normal distribution with a mean different from zero. AS is characterized by a platykurtic or bimodal distribution with a mean of zero. Most DA corrections essentially consist in considering FA as deviations around the mean signed right-left asymmetry (mean DA) in the sample instead of as deviations around zero. For example, mean DA can be subtracted from the individual asymmetry values, or corrected for by ANOVA, using the fixed side effect to quantify it.

One main purpose of geometric morphometrics (GM) is to quantify shape information and analyse it in subsequent mathematical procedure. Once the landmarks are taken, Procrustes superimposition is applied. This superimposition takes away three redundant information, scale, position, and rotation. Scale is often eliminated by setting the centroid size, square root of sum of squared distances between the centroid and each landmark, the same in all specimens.

“Companion” rabbits resemble juvenile stages: large eyes in relation to face size, a large head disproportional to the body.) They can be considered as paedomorphic. Pet product marketers, certainly have been taking advantage of these phenomena and their implications, and recently they have created many companion rabbit breeds, inducing shifts in their development for getting the “cute factor”. Certainly, human mental model to both types of animals respond to the same “cuteness” aspect. They can be considered paedomorphic. Paedomorphosis refers to underdevelopment, so that the adult passes through fewer growth stages and resembles a juvenile stage of its ancestor (4). It results in a reduction in the rate of developmental changes (5), requiring less growth of highly developed adult body forms (6). It appears either when character development is delayed or through acceleration of sexual maturation (7). A large head and a round face, a high and protruding forehead, large and low-lying eyes, bulging cheeks, and a small nose and mouth are some of the components of this quality (8). This “babyness” is perceived as attractive and cute by humans (8). Studies have shown that humans find paedomorphic features more attractive. This is mostly put down to the infantile features tickling a subconscious need to care for a younger individual, including animals (8). Physical paedomorphism has been described in domestic dogs (9), which is characterized by a reduction in overall body size and retention of a juvenile head:body ratio (in (9)). It has also been cited in cats (10) and horses (9), but, at least to the authors knowledge, nothing has been published in rabbits.

To fill this gap we report here preliminary findings that for the first time to quantify size asymmetry mandibles of companion rabbits. More specifically, we addressed the following questions: (1) Which types of mandibular size asymmetries occur in companion rabbit? (2) Does the level of detected asymmetries vary according to body size? (3) Can we infer that the extreme selective traits in companion rabbits create abnormal functional conditions, which in turn could be expressed as high degree of asymmetry?

## Materials and methods

### Sample

A total sample of 64 freshly dead companion rabbits (17 males and 46 females, and one of unregistered gender) was studied. They were from the same farm, and managed in identical conditions (housing, feeding, preventive treatments…). At the laboratory room, corpses were sexed and weighed (range 0.27-3.37 kg, mean weight 1.50±0.59 kg) and the heads were excised. The defleshing process was done naturally using scavenging beetles and flies. Once completed, the heads were thoroughly washed in water and allowed to dry at room temperature. Mandibles were then extracted, macerated with water and finally whitened with hydrogen peroxide.

All mandibles are currently deposited on the collection of the Department of Animal Science of the University of Lleida, and more information can be sent upon request to the first author.

### Photographs and landmark data

Digital photographs of right disarticulated hemimandible on their lateral aspect were obtained. Digital capture was performed with a Nikon® D70 digital camera (Nikon Inc., Tokyo, Japan) (image resolution 2,240 x 1,488 pixels) equipped with a Nikon AF Nikkor® (Nikon Inc., Tokyo, Japan) 28 to 200-mm telephoto lens. Imaging procedure were standardised as follow: hemimandibles were set as to rest on their medial side, the focal axis of the camera being parallel to the lateral aspect. The camera was attached to a column with an adjustable arm, and above a grid baseboard for measure reference.

The mandible morphology was described by a set of 18 landmarks and semilandmarks (Figure 1) covering the body and the ramus, and assumed to be homologous and topologically equivalent. Landmarks used in this study were primarily chosen to describe major mandibular regions as well as parts of particular morphofunctional interest. They refer to the: 1) distal edge of the incisive alveolus; 2) medium part of the diastema; 3) mental foramen (most caudal edge); 4) first premolar alveolus (most oral point); 5) last molar alveolus (most caudal point); 6) half part of the *processus coronoides;* 7) deeper part of the *incisura mandibulae;* 8) most rostral part of the *caput mandibulae;* 9) most caudal part of the *caput mandibulae;* 10) deeper part of the *ramus mandibulae;* 11) angular process; 12) half edge of the *fossa masseterica;* 13) most ventral angle of the *fossa masseterica;* 14) mandibular notch; 15) ventral projection of 5); 16) ventral projection of 4); 17) ventral projection of 2); and 18) incisor alveolus (ventral). The choice of alveolar rather that tooth crown landmarks was motivated by the fact that some of our specimens lacked some of these teeth. All these points were chosen according to their potential accuracy of digitization and because points were homologous through the structures, furthermore they would represent the mandible and its parts as good as possible: the mandible body (*corpus mandibulae,* horizontal part, landmarks 1 to 5, and 15 to 18, these latter utilizing perpendicular projected points on the ventral border in relation to dental position, and 1, 4 and 5 recorded at the alveolar edges adjacent to the teeth) and ramus *(ramus mandibulae,* vertical part, landmarks 6 to 14). The chosen landmark configurations occupy different regions of the theoretical morphology defined by mandibular apparatus (11). No differences according to coat were considered.

**Figure 1.**
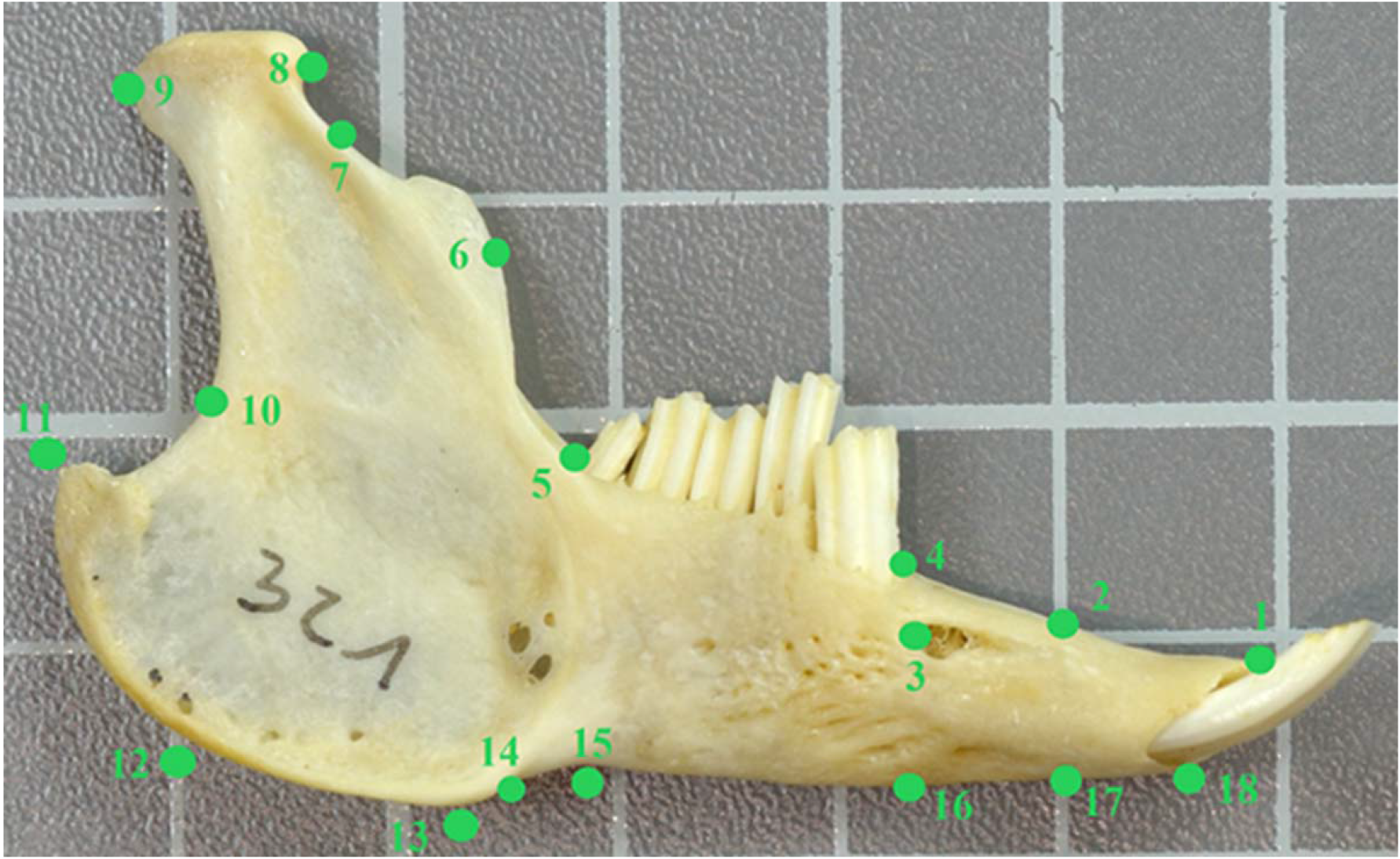
Picture illustrating a set of recorded landmarks and semilandmarks used. Pictures were taken on the lateral aspect of both hemimandibles. See text for a detailed anatomical description of each landmark.

Each landmark was digitalized two times independently in each two hemimandibles by the two authors to allow for estimating replicability. The Cartesian coordinates of all landmarks were digitized using TpsDig, v. 1.40 software (12). Replicability of Procrustes coordinates was then analysed by a two-way NPMANOVA (Non-Parametric-Multivariate-Analysis-Of-Variance) using 9,999 permutations and Euclidean distances with sides (S) and replicas (R) as factors. For shape asymmetry, configurations superimposed were used as dependent variables, such that the effect of the side corresponded to directional DA, the interaction between the side of the body (S) and the replica (R) corresponded to FA, and the residual term corresponded to the measurement error in the model (3). In fact, the ratio of the R-by-S mean square to the combined R-by-S-by-R and S-by-R mean squares provided an *F*-test of whether between-individual variation in estimated asymmetry can be accounted for by measurement error. This set was further standardized by the Generalized Procrustes Analysis (GPA). GPA begins by reflecting landmark configurations from one of the sides and superimposing them by their centroid (midpoint of a configuration of anatomical landmarks). The size of the centroid (CS) is a side product of the GPA fitting and is computed as the average distance between landmarks and the centre of gravity of a given configuration. In this context is defined as the information that remains in a set of coordinates after these parameters have been removed. The CS data can be analysed similarly to asymmetries of ordinary metric traits. Finally, each landmark configuration was rotated such that the squared distances between homologous landmarks were minimized.

As a result of all of these calculations, CS from averaged Procrustes coordinates was obtained. As non-normal distribution appeared (Shapiro-Wilk’s *W*=0.283, *p*⋘0.001), symmetry was studied by means of non-parametric tests, Kolmogorov-Smirnov *D*, Mann-Whitney *U* and Wilcoxon paired *W*. The validity of FA interpretations depends on an absence of DA, AS and a normal distribution for right minus left with mean zero (13). To reinforce whether data conformed to the requirements for a trait showing FA, we used Mann-Whitney tests to determine if the relative measures of asymmetry differed significantly from zero. As males and females presented an overall equal distribution (*D*=0.127, *p*=0.762), sex was not a prior consideration. As relative measures of asymmetry, we used signed right minus left CS differences (14) were obtained for distribution study. Size dependence of FA (to correct for possible associations between asymmetry and size) was evaluated within using the significance of the Spearman Rank correlation coefficients of absolute asymmetry ([right-left]) on character size defined as ([right+left]/2) (15). To measure the direction and magnitude of asymmetry we used the percentage of DA by calculating the difference between a left and right pair of measurements, standardized by the mean of the left and right measurements [(R-L)/ {(R+L)/2)}] × 100%. This calculation was chosen because it offers a different way of expressing DA, as it eliminates potential problems associated by the use of descriptive statistics in calculating %DA (%DA is a signed number and can lead to the generation of mean and standard deviation values that do not reflect the true differences). Percent bias [{Count (R>L)/ Count (R)}] × 100% was used as a means of calculating asymmetry as a count variable. Finally, a simple linear regression of body weight (data log-transformed) with signed differences were obtained. In this case, the Wilks′ *λ* test statistic was computed as the ratio of determinant. All analyses were carried out using the softwares MorphoJ v. 1.06c (16) and PAST v. 2.17c (17).

### Ethics Statement

This study was carried out in corpses from naturally dead animals by causes other than the purpose of this study so no Ethics Committee agreement was considered.

## Results

Replicas were shown to be highly repeatable indicating a very low influence of error on measurements. In other words, the side variation in estimated asymmetry was significantly larger than within-side variation due to measurement error. The value of absolute asymmetry was independent of size of the trait (r_s_=0.026, *p*=0.832). Therefore, we did not correct asymmetry measures for a size-dependent relationship (table 1).

**Table 1.** Two-way NPMANOVA (Non-Parametric-Multivariate-Analysis-Of-Variance), using 9,999 permutations and Euclidean distances for hemimandibular raw coordinates, with sides and replicas as factors, for 64 companion rabbits. The individual amount of variation for sides exceeded the digitalization error suggesting than for this study, digitalization error was not a concern.

| Source | Sum of squares | Degrees of Freedom | Mean square | *F* | *p* |
| --- | --- | --- | --- | --- | --- |
| Side | 2.35E+08 | 1 | 2.35E+08 | 208.24 | 0.0001 |
| Replica | 1.97E+06 | 1 | 1.97E+06 | 1.7442 | 0.1761 |
| Interaction | 4.04E+05 | 1 | 4.04E+05 | 0.35838 | 0.7361 |
| Residual | 2.84E+08 | 252 | 1.13E+06 |  |  |
| Total | 5.21E+08 | 255 |  |  |  |

Mann-Whitney *U* reflected no differences between sides (*U*=1881, *p*=0.427), but Wilcoxon paired test reflected differences (*W*=1678, *p*⋘0.001). Normality tests indicated that hemimandibles showed evidence of DA too (*U*=1024, *p*⋘0.001), asymmetry distributions thus being non normal with mean –5.2, neither bimodal (left skew of −1.890) nor leptokurtic (kurtosis of 6.658) (figure 2). No significant regression appeared with body weight (data log-transformed) (*R*^2^=0.028, Wilk’s *λ*=0.971, *F*_1,62_=1.811, *p*=0.183) (figure 3). Overall, the mean DA represented −1.0%±1.87 of trait size and percent bias was 25.0%.

**Figure 2.**
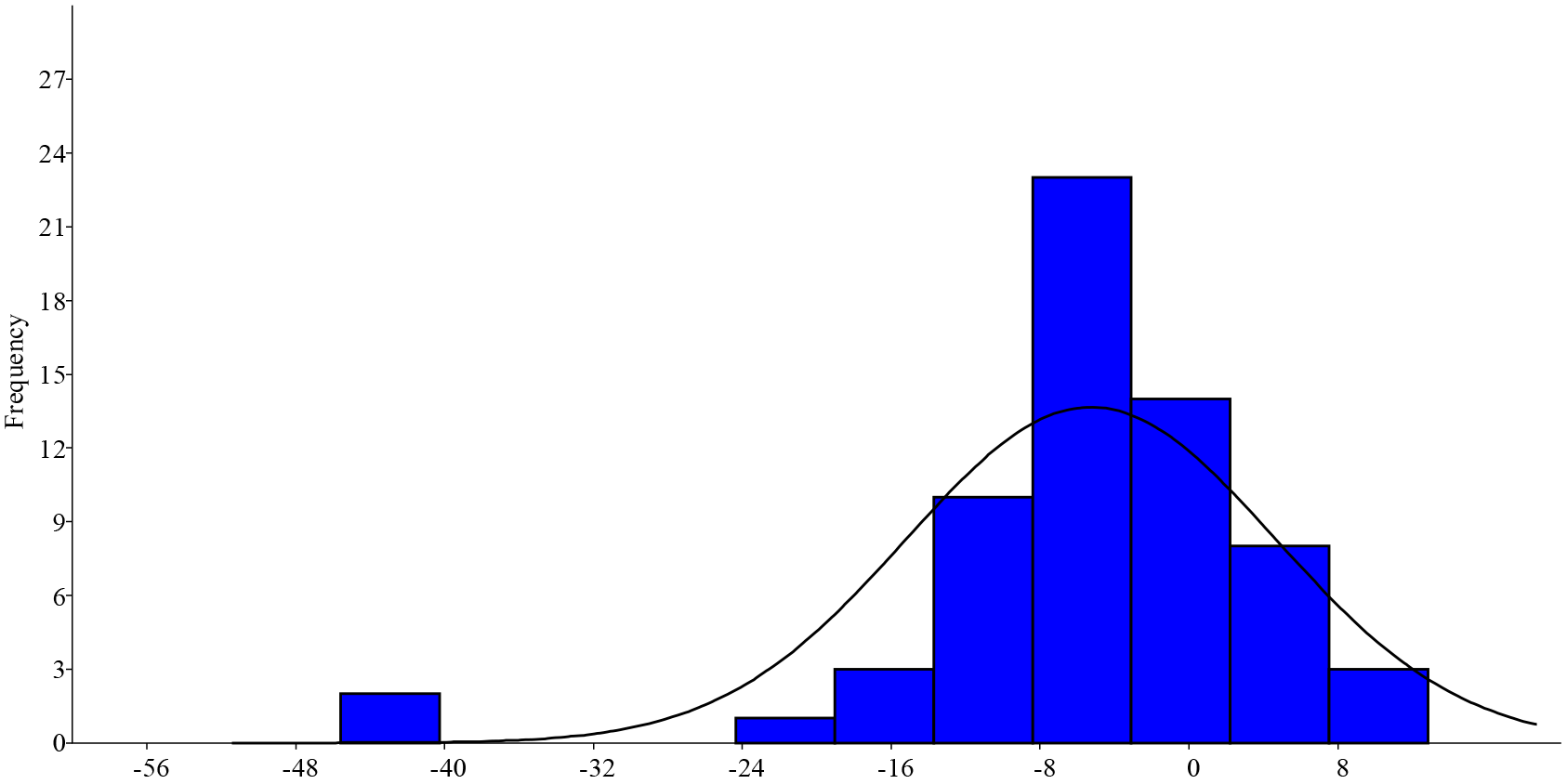
Distribution of signed right-left CS differences for 64 companion rabbits. Distributions was non normal with mean −5.2, being neither bimodal (left skew of −1.890) nor leptokurtic (kurtosis of 6.658).

**Figure 3.**
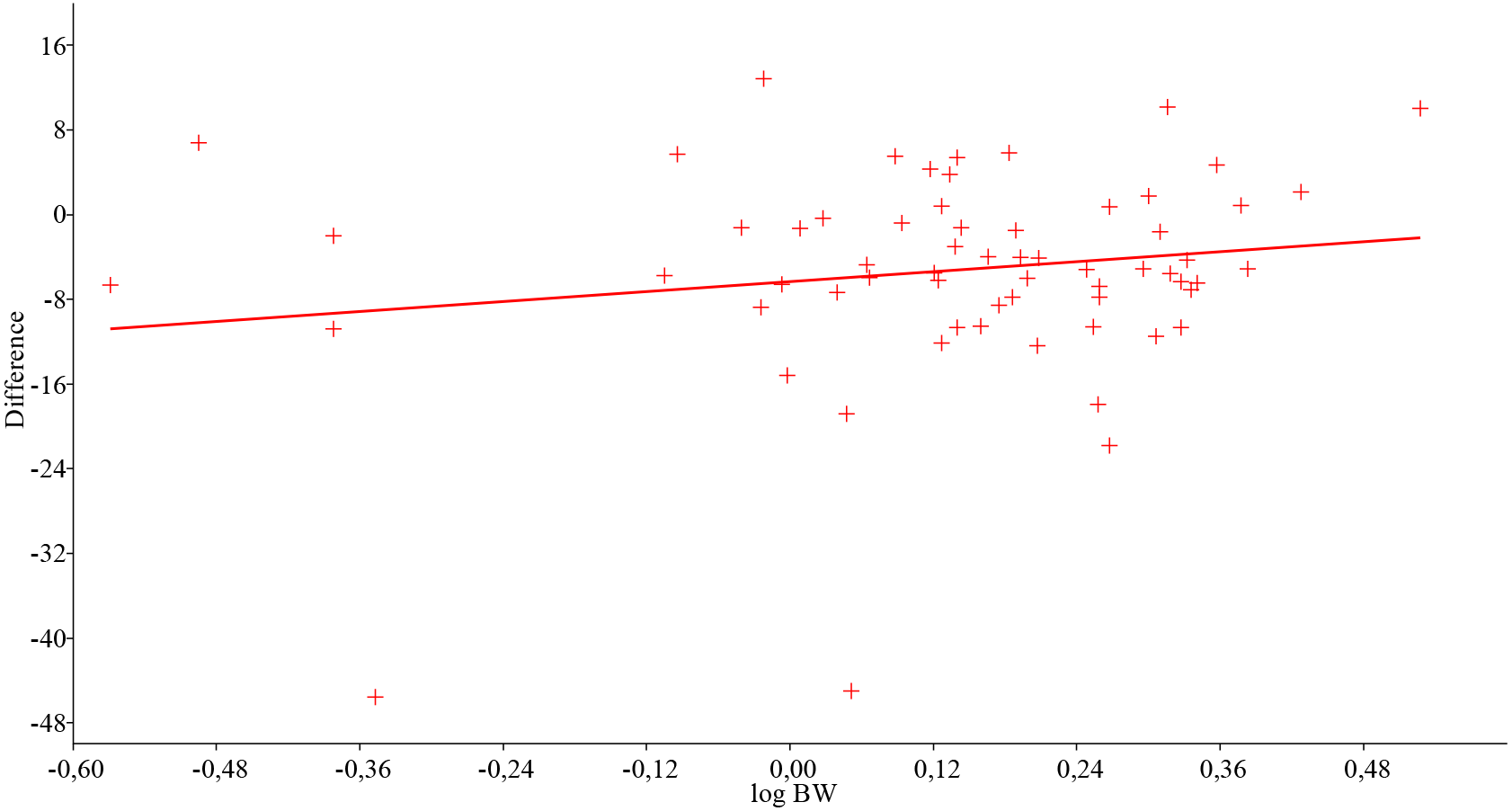
Regression of body weight (BW, data log-transformed) with signed right-left centroid size differences, for 64 companion rabbits. No significant regression appeared with body weight (*R*^2^=0.028, Wilk’s *λ*=0.971, *F*_1,62_=1.811, *p*=0.183).

## Discussion

DA refers to significant unimodal population-level deviations from bilateral symmetry that most likely arises from lateralized behaviours, and in the present cases it seems it should not indicate an index of DI. It is noteworthy that the standard deviation is greater than the mean value. This is due to the fact that % DA is a signed value, and, because of bias in its calculation, it may not represent the true mean of the population. Using % Bias as a measure of asymmetry avoids this problem. It had values of 25%; this value indicates a non-significant right bias.

In general, for companion rabbits, our detected DA would favour left mandibular side. This unilaterality could be interpreted as a manifestation of lateralized masticatory activity, the main explanation for the detected mandible’s asymmetry is a biomechanical mandibular laterality. Other possible factors such as genetic and hormonal development focused their explanations of the differences on vascular supply and environmental stress factors; as malnutrition and extreme weather which seems unlikely at least under the current management system of the studied animals, with high welfare standards.

Bone is a dynamic tissue, which continuously undergoes adaptive remodelling, i.e. resorption and apposition, to meet the requirements of its functional environment (18). Laterality addresses asymmetry in size of mandibles by way of mandible remodelling rate, including depository and resorptive processes. For instance, on the ramus, the lingual surface is predominantly depository, in contrast to the resorptive nature of the contralateral buccal side (19). Remodelling differences in mandibular body are present, too (19). Under physiological conditions, intermittent mechanical loading of bone is caused predominantly by muscle recruitment and contractions (18). The muscles thus provide an important mechanical stimulus for bone remodelling by inducing strains in the skeletal system (18). This remodelling would be gross enough to express size differences, and perhaps more linked to food hardness (a reduction in mechanical loading of the mandible brought about by mastication of soft food has been said to decrease the remodelling rate of bone, which, in turn, might increase the degree of bone mineralization (18). As mechanical loading during mastication is not evenly distributed over the mandible, this increase might be regionally different (18). A further study of masticatory muscle mass in relation to mandible bone shape will further elucidate the contributions of muscle development, contraction and biomechanical loading in lagomorphs.

## Conflicts of interest

The authors declare no conflicts of interest.

## Acknowledgement

Authors thank CUNIPIC, in Vallfogona de Balaguer for offering cadavers of their animals. We are very grateful to anonymous referees for their useful suggestions.

